# Proteoform Barcode: An Intuitive Visualization Framework for Top-Down Proteomics

**DOI:** 10.64898/2026.08.17.745297

**Authors:** Yifan Yue, Guangyao Gao, Fei Fang, Guijie Zhu, Seyed Amirhossein Sadeghi, Reyhane Tabatabaeian Nimavard, Liangliang Sun

**Author notes:** Corresponding authors: Liangliang Sun.

## Abstract

Top-down proteomics (TDP) advances biomedical research by providing a bird’s-eye view of proteoforms in cells, tissues, and biofluids. Thousands of proteoforms can be characterized using well-established TDP technologies, and potential proteoform biomarkers of diseases have been discovered. However, there is a lack of an easy and biologically informative approach to present the quantitative global TDP data. Here, we present “proteoform barcode” as a straightforward visualization approach that simultaneously displays proteoform abundance and their associated Gene Ontology (GO) biological processes, converting a list of proteoforms to a biologically informative image. The proteoform barcode allows 1) a global view of proteoforms (i.e., relative abundance and functional information) in complex biological systems (i.e., bacteria, yeast, human cells, and human plasma) and 2) the accurate distinction of samples in diverse biological conditions (i.e., control and disease) assisted by machine learning approaches. The proteoform barcode, assisted by the random forest model, accurately separated the human plasma samples of healthy controls and early-stage breast cancer. The data demonstrates the high potential of the proteoform barcode-based approach for early diagnosis of diseases in an easy and biologically informative manner.

**For TOC only:** 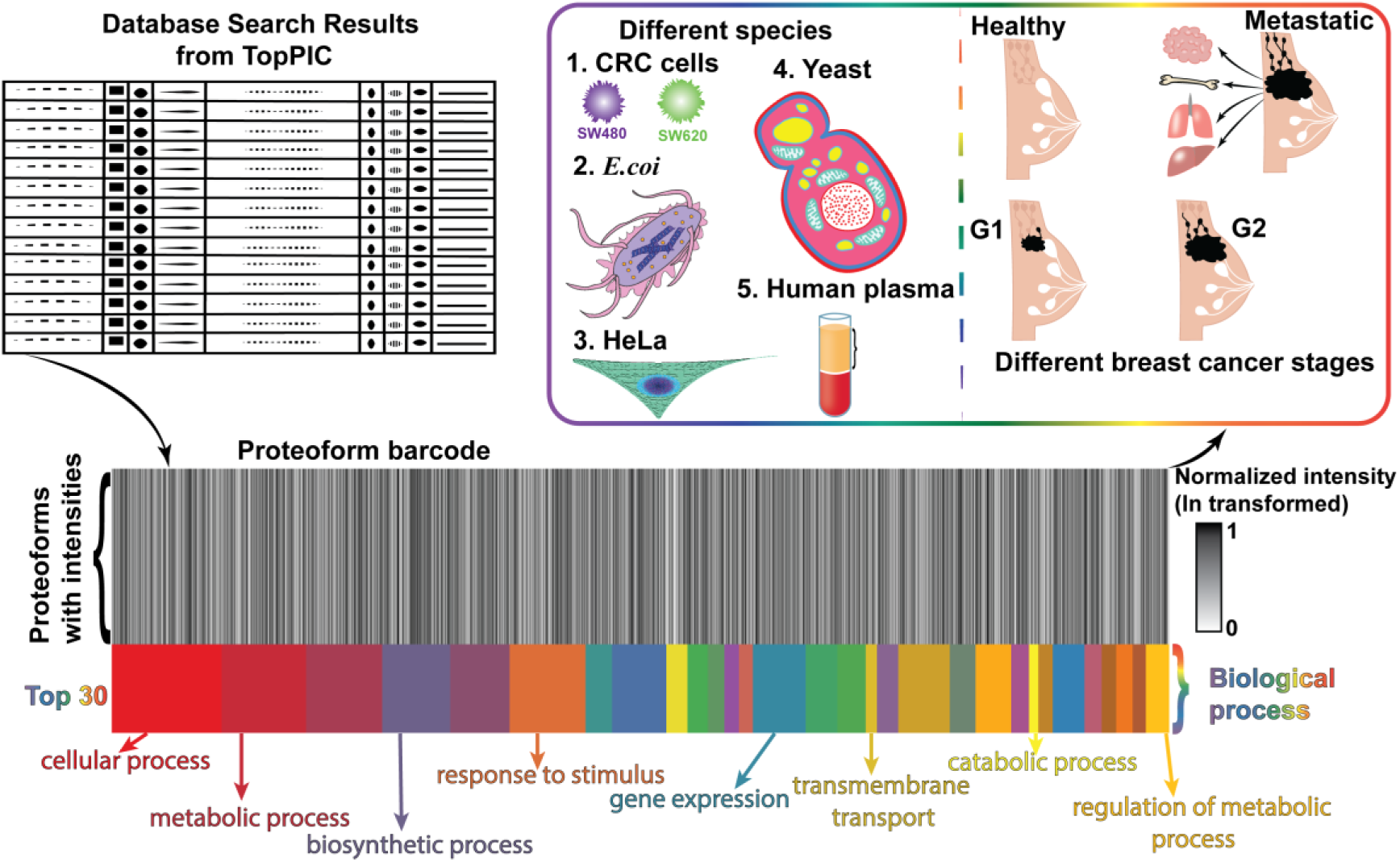

## 1. Introduction

Top-down proteomics (TDP) has evolved over more than three decades, driven by the development of soft ionization techniques (i.e., electrospray ionization, ESI)^1^, high-resolution mass analyzers (i.e., Orbitrap)^2^, highly efficient gas-phase fragmentation techniques [i.e., higher-energy collisional dissociation (HCD)^3^, electron transfer/capture dissociation (ETD/ECD)^4–6^, and ultraviolet photodissociation (UVPD)^7, 8^], advanced sample preparation techniques^9^, high-resolution chromatographic or electrophoretic separations^10, 11^, and bioinformatics tools^12–16^ for database search-based proteoform identification and relative quantification. Thousands of proteoforms can be identified and/or quantified using well-established TDP technologies in complex biological samples^17–21^, including human^22–35^, zebrafish^36–39^, mouse^40–44^, and *Escherichia coli (E. coli)* ^36, 45–47^. A large number of TDP datasets have been generated in the last 10 years, making an easy and biologically informative approach for data visualization critical to fully use the available TDP data.

Typically, the TDP studies produce a long list of identified proteoforms or differentially expressed proteoforms between biological conditions. Additional sophisticated data analyses are needed to gain biologically informative information from the TDP data. For example, it is crucial to connect the identified and quantified proteoforms to the biological processes or pathways that they are involved in by Gene Ontology (GO) analysis. For another example, a group of differentially expressed proteoforms between disease and healthy control can be used as biomarkers of disease. Usually, machine learning or artificial intelligence (AI) tools are needed to improve the accuracy of disease diagnosis using the proteoform biomarkers. A simple and biologically informative data presentation approach for the differentially expressed proteoforms will facilitate machine learning or AI-based disease diagnosis.

To accelerate progress in using TDP data for disease diagnosis and facilitate the extraction of biologically informative information from the TDP data, we developed the “proteoform barcode” as an intuitive visualization framework for TDP. In this visualization, the x-axis represents proteoforms grouped and ordered by their associated biological processes. Each vertical line corresponds to an individual proteoform, with its color indicating the normalized log-transformed intensity. The colored bands at the bottom denote different biological processes, enabling researchers to quickly assess proteoform abundance patterns and biological process-level changes across samples. We then compared proteoform barcodes across different species and biological conditions. Remarkably, proteoform barcode images generated from human plasma samples of controls and patients with stage G1 breast cancer were used for the diagnosis of early-stage breast cancer, assisted by a machine learning approach.

## 2. Methods

### 2.1. The procedure for producing proteoform barcodes

The proteoform barcode generation is summarized in **Figure 1**, which includes seven main steps. **The first step is data collection.** All datasets used in this study were obtained from previously published studies. The dataset 1 contains proteoforms identified from metastatic (SW620) and nonmetastatic (SW480) colorectal cancer (CRC) cell lines^22^. Dataset 2 contains *E. coli* proteoforms from ref.^47^. Dataset 3 contains HeLa cell proteoforms from our previous work^48^. Dataset 4 contains yeast cell proteoforms from ref.^49^. Dataset 5 contains human plasma proteoforms from 39 samples representing four stages of breast cancer (9 Controls, 10 G1, 10 G2, and 10 metastatic stages)^29^. Datasets 1 and 2 are directly obtained from the online supplementary information of the original publications. Datasets 3–5 are available at https://github.com/YiFanYUE99/proteoform_barcode. For all the database searches, TopPIC (TOP-down mass spectrometry-based Proteoform Identification and Characterization) was used^12^.

**Figure 1.**
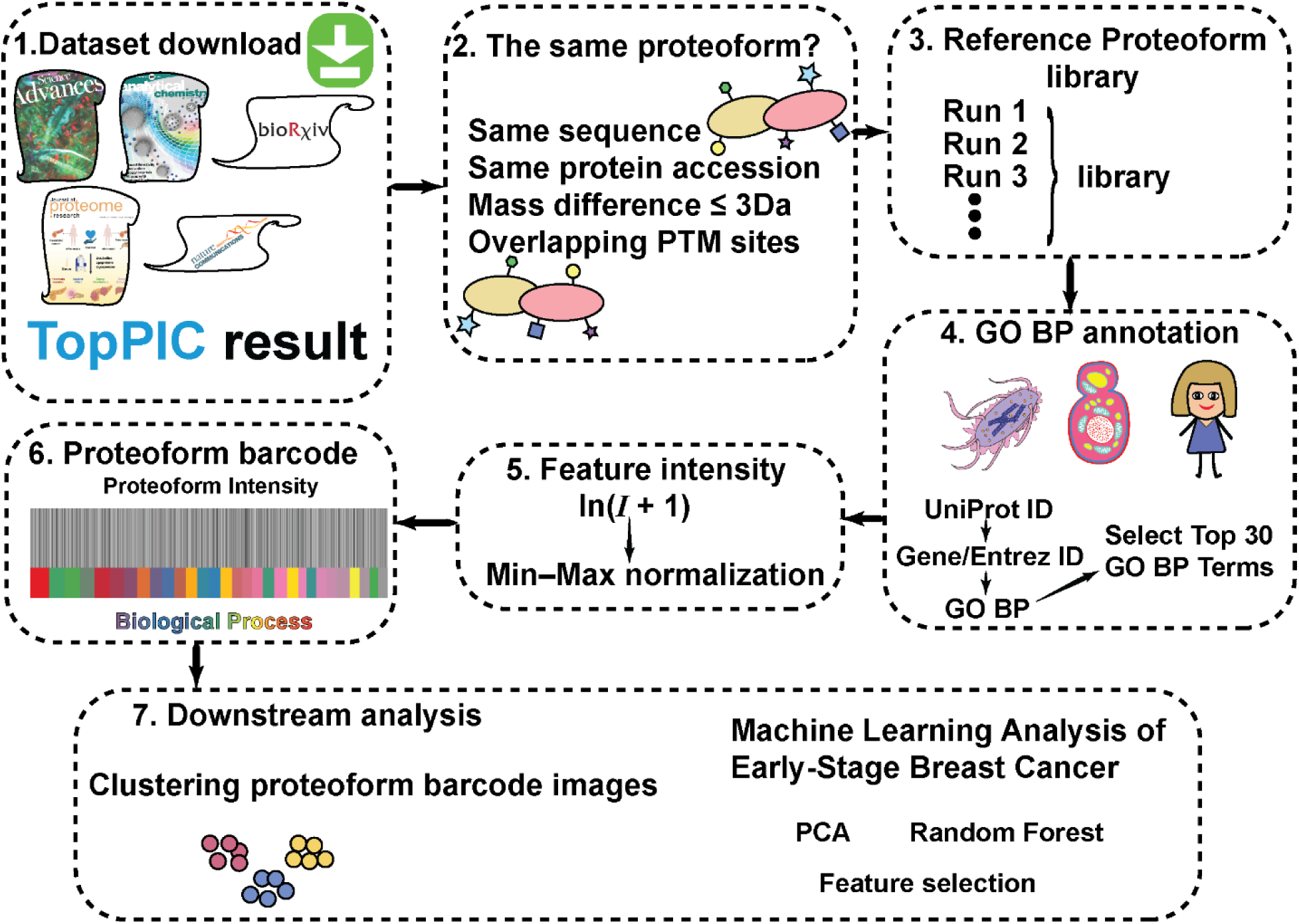
Workflow for proteoform barcode generation, clustering, and its potential application in early disease detection.

#### In the second step, we tried to determine whether proteoforms are the same or not

Proteoforms were considered identical only if they met all of the following criteria: (1) the same protein sequence, (2) the same protein accession number (i.e., same UniProt accession number), (3) a mass difference of no more than 3 Da, and (4) overlapping post-translational modification (PTM) sites, when PTMs were present. As shown in **Table S1**, proteoforms 1 and 2 share the same protein sequence and protein accession, and their mass difference is less than 3 Da. However, they are not considered the same proteoform because their PTM sites do not overlap. In contrast, proteoforms 3 and 4 are classified as the same proteoform because they have the same protein sequence, the same protein accession, a mass difference of less than 3 Da, and overlapping PTM sites.

#### The third step is data preprocessing before making the proteoform barcode

All proteoform barcodes presented in this study were generated from the database search results of the TopPIC^12^. The input can be either a TSV file generated from a single MS run or a combined TSV file from multiple MS runs. The following columns from the TopPIC output were required for proteoform barcode generation: Adjusted precursor mass, Proteoform, Protein accession, and Feature intensity. In older versions of TopPIC, or when the database protein sequence was not included in the exported database search results, the database protein sequence was extracted directly from the Proteoform column. Specifically, all characters preceding the first period (.), including the period itself, and all characters following the last period, including the period, were removed. In addition, modification annotations enclosed in square brackets ([]), parentheses, and hyphens (-) were removed, yielding the database protein sequence for downstream analysis.

#### In the fourth step, reference proteoform libraries need to be generated

After proteoforms identified from all MS runs within a project were grouped according to the criteria described in the second step, each unique proteoform was assigned to a unique ‘cluster_id_with_mod’. Within each ‘cluster_id_with_mod’, only the proteoform with the highest feature intensity was retained as the representative proteoform, while all other entries were removed. A reference proteoform library was then generated containing the Database protein sequence, Adjusted precursor mass, Proteoform, Protein accession, and ‘cluster_id_with_mod’. The reference proteoform libraries of different datasets are in **Supporting Information II** (**Sheet S1-S6**).

#### In the fifth step, proteoforms were connected to their GO Biological Processes and top 30 GO terms were chosen

For human, yeast, and *E. coli*, GO annotations were retrieved from ‘org.Hs.eg.db’, ‘org.Sc.sgd.db’, and ‘org.EcK12.eg.db’databases, respectively, using Entrez Gene IDs as the key type. All genes in each organism database were extracted and mapped to their associated GO annotations. Only annotations belonging to the BP were retained. To avoid counting the same GO term multiple times for the same gene, duplicated genes were removed. The top 30 GO Biological Process (BP) terms were selected separately for each organism using the corresponding organism-specific annotation database.

The number of unique genes associated with each GO BP term was then counted, and GO IDs were converted to readable GO term names using ‘GO.db’. GO BP terms were ranked in decreasing order according to the number of associated genes, and the top 30 most abundant BP terms were selected for each organism. A unified color map was generated across all selected GO BP IDs so that the same GO BP term was assigned the same color across different organisms (human, yeast, and *E. coli*). The top 30 BP for the three organisms are provided in the **Supporting Information II** (**Sheets S7–S9**).

#### In the sixth step, GO BP annotations were performed

GO BP annotations were added to the proteoform library by extracting UniProt IDs from the Protein accession column and mapping them to Entrez Gene IDs using ‘org.Hs.eg.db’(for human samples) and ‘org.Sc.sgd.db’(for yeast samples) annotation databases. Entrez IDs were then used to retrieve associated GO annotations, and only BP terms were retained. For the *E. coli* dataset, GO annotations were retrieved using the mapped gene names because the corresponding Bioconductor annotation database is no longer maintained. UniProt accessions extracted from the Protein accession column were mapped to gene names using the UniProt ID Mapping service (https://www.uniprot.org/id-mapping), and the resulting gene names were subsequently used for GO BP annotation. The resulting GO BP annotations were merged back into the proteoform library based on UniProt ID. Finally, only proteoforms annotated to the selected top 30 GO BP terms were retained for barcode construction. The selected BP terms were arranged from left to right in descending order according to the number of annotated genes in each BP, providing a consistent organization of proteoform barcodes across all samples within the same dataset.

#### In the seventh step, intensity log transformation and normalization were performed for constructing a single barcode from all runs of the same sample

For constructing a single barcode by integrating multiple MS runs from the same biological sample or species, feature intensities were processed independently within each dataset. When the same proteoform (defined by the same cluster_id_with_mod) was identified multiple times, either across different runs or multiple times within the same run, only the highest feature intensity was retained for subsequent analysis. A natural logarithm transformation [ln(*I* + 1)] was first applied to the feature intensities. The log-transformed intensities were subsequently normalized to the range of 0–1 using min–max normalization according to the following equation:

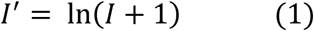

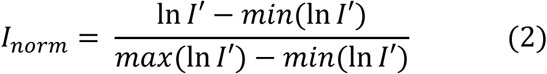

where *I* represents the feature intensity. This normalization was performed independently for each dataset and was used solely for barcode visualization when integrating multiple MS runs into a single barcode.

#### Finally, proteoform barcode was generated

Proteoform barcodes were generated in R using the *ggplot2*^50^ package based on the annotated proteoform library containing only proteoforms assigned to the selected top 30 BP terms. Each barcode represented the integrated proteoform profile from all runs of the same biological sample or species. Proteoforms were displayed according to their assigned BP terms, with BP categories arranged in descending order according to the number of unique genes annotated to each term. Proteoforms associated with multiple BP terms were displayed within each corresponding BP category. The upper portion of each barcode represents normalized proteoform intensities. Each proteoform was plotted as a single vertical line, with the line color corresponding to its normalized intensity (0 = white and 1 = black). The lower portion of the barcode represents BP annotations, in which each proteoform was displayed as a colored rectangle corresponding to its assigned BP term. The same color was consistently assigned to the same BP term across all barcodes.

All barcodes were exported as PDF files with a fixed size of 8 inches in width and 2 inches in height to ensure consistent dimensions across all datasets. The PDF files were subsequently converted to PNG format using Adobe Acrobat to preserve the original BP colors and prevent color distortion during image conversion.

### 2.2 Proteoform barcode generation for breast cancer stage comparison (Dataset 5)

For the breast cancer dataset (**Dataset 5**), feature intensities were first organized at the individual technical run level. When the same proteoform (defined by the same cluster_id_with_mod) was identified multiple times within a technical run, only the highest feature intensity was retained. An intensity matrix was then generated in which each technical replicate run was represented by a separate column, with missing proteoform detections assigned a value of zero.

Each biological replicate consisted of three technical replicate runs. To obtain a representative intensity for each biological replicate, the mean feature intensity of each proteoform was calculated across the three technical replicates using only detected (nonzero) intensities. If a proteoform was not detected in any of the three technical replicates, its intensity was assigned a value of zero. The resulting biological replicate-level intensities were then natural log-transformed [ln(*I* + 1)] and normalized according to Equations (1) (2) before barcode construction and downstream comparison of different breast cancer stages.

Proteoform barcode generation was performed as described above using the annotated proteoform library containing only proteoforms assigned to the selected top 30 BP terms. Unlike the integrated barcode generated from multiple MS runs, each barcode represented a single biological replicate. All other barcode generation procedures, including proteoform ordering, color mapping, image dimensions, and PDF to PNG conversion, were identical to those described above.

### 2.3 Clustering different breast cancer stages using the proteoform barcodes

PNG barcode images were resized to 800 × 200 pixels. Each image was converted to RGB format, and pixel values were inverted so that white background pixels contributed minimally while darker or colored barcode pixels contributed larger values. For each image, red, green, and blue channel intensity profiles were calculated by summing pixel intensities column-wise across the image height. The three channel profiles were concatenated into one feature vector. Each feature vector was normalized by Z-score transformation to reduce the influence of overall image brightness. The resulting feature matrix was further standardized across samples. Hierarchical clustering was then performed directly on the standardized RGB profile matrix using correlation distance and average linkage. For clustering involving two stages, the number of clusters was set to two. When all samples were clustered simultaneously, the number of clusters was set to four.

### 2.4 Multivariate analysis and classification of different biological conditions

PCA was performed using the R function prcomp on the log-transformed and normalized breast cancer proteoform dataset without centering or scaling (center = FALSE, scale. = FALSE) to visualize sample distribution. A random forest classifier was implemented using the scikit-learn library to distinguish Control and G1 samples, with the aim of evaluating the potential of proteoform barcodes for early breast cancer diagnosis. The dataset was randomly split into training and test sets with a test size of 40%. The classifier was trained using 500 decision trees with a fixed random seed (random_state = 66). Model performance was further evaluated using stratified 5-fold cross-validation with shuffling enabled, using the same random seed, and receiver operating characteristic (ROC) curves were generated to assess classification performance. The top 100 most important proteoforms were used to construct the heat map and optimized proteoform barcodes with improved visual discrimination between Control and G1 samples.

### 2.5 Proteoform barcodes generated using the top 100 proteoform features for early breast cancer diagnosis

To better visualize the differences between Controls and patients with stage G1 breast cancer, new proteoform barcodes were generated using the top 100 proteoforms selected by the random forest model above. All these 100 proteoforms were associated with “cellular process”, which is the top 1 GO BP in humans. For barcode construction, only cellular process GO BP were retained, and annotations to other biological processes were ignored.

## 3. Results and Discussion

### 3.1 Proteoform barcode reflects the proteoform profile differences across different biological samples

A summary of the reference proteoform libraries for all datasets is provided in **Table 1**, including the total number of identified proteoforms and the corresponding proteoforms retained after filtering by the selected top 30 GO BP terms for barcode construction. The percentage of proteoforms retained within the selected top 30 GO BP terms ranged from 80% to 99% across all datasets. These results suggest that using only proteoforms associated with the selected top 30 GO BP terms largely preserves the information contained in the original proteoform libraries.

**Table 1.** Summary of proteoforms in each reference proteoform library and the corresponding proteoforms retained after filtering by the selected top 30 GO BP terms for proteoform barcode.

| Dataset | Species | Raw files | Total Proteoforms | Proteoforms in Top 30 BPs | Percentage |
| --- | --- | --- | --- | --- | --- |
| CRC cells (SW480) | Human | 214 | 17065 | 16102 | 94% |
| CRC cells (SW620) | Human | 198 | 14334 | 13580 | 95% |
| *E. coli* | *E. coli* | 43 | 4554 | 3624 | 80% |
| HeLa cells | Human | 31 | 4697 | 4118 | 88% |
| Yeast | Yeast | 62 | 10805 | 10637 | 98% |
| Human plasma | Human | 27 | 2087 | 2057 | 99% |
| Breast cancer plasma dataset | Human | 117 | 3845 | 3785 | 98% |

To evaluate the general applicability of the proposed proteoform barcode strategy, six proteoform barcodes are created from five representative datasets, including CRC cell lines (SW480 and SW620), *E. coli*, HeLa cells, yeast, and human plasma, **Figure 2**. In each barcode, every vertical line represents a proteoform, with grayscale intensity indicating its normalized intensity. Proteoforms are arranged according to their assigned GO BP terms, and the colored band beneath the barcode indicates the corresponding BP category. The proteoform barcode therefore integrates quantitative proteoform intensity with biological process annotation into a single intuitive visualization.

**Figure 2.**
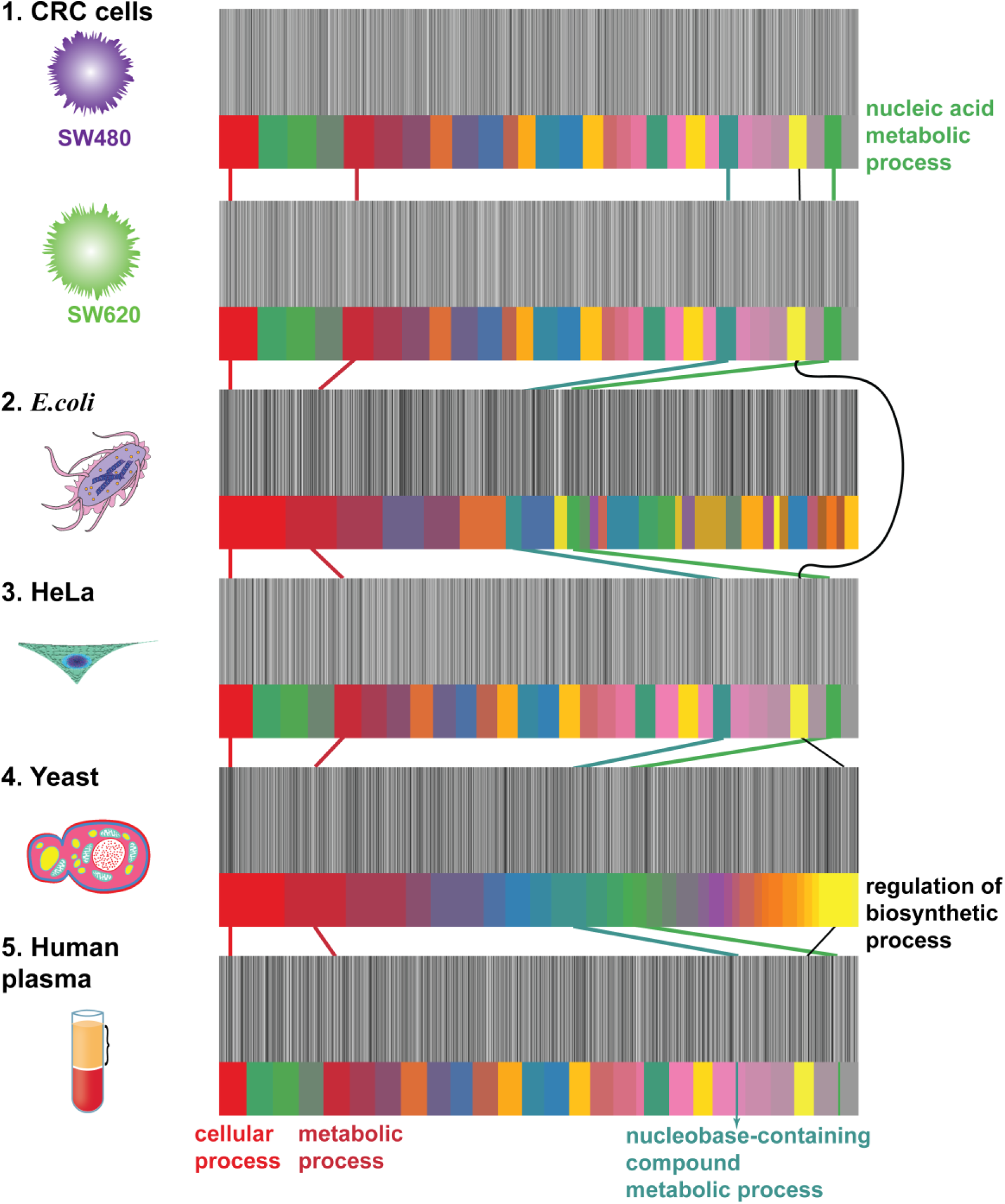
Proteoform barcodes from different sample types and species. From top to bottom: CRC cells, E. coli, HeLa cells, yeast, and human plasma. Several example biological processes (BPs) were annotated and connected across sample types by color-matched solid lines. Each colored band at the bottom represents a different biological process (BP) and contains multiple vertical lines, with each vertical line representing a proteoform associated with that BP. Because a proteoform may be associated with multiple BPs, the same proteoform may appear in multiple colored bands. The grayscale gradient of the vertical lines represents normalized intensity of each proteoform, ranging from 0 (white) to 1 (black); the same scale is used in all subsequent figures.

Although the CRC cell lines, HeLa cells, and human plasma datasets were all annotated using the same human GO BP database, they exhibited distinct color band patterns, indicating different distributions of proteoforms among different BP categories. The data indicates the potential of using specific proteoform barcodes to represent different human samples, i.e., different cell types. Cellular process and metabolic process were present in all three species and occupied a larger proportion of the proteoform barcodes in yeast and *E. coli* than in the human datasets. Regulation of biosynthetic process was absent in *E. coli* but present in the other species. According to the yeast GO annotation database, this BP contains the fewest annotated genes among the selected top 30 BP terms. Nevertheless, it still occupied a noticeable region of the yeast proteoform barcode. It is also worth noting that nucleic acid metabolic process and nucleobase-containing compound metabolic process were well represented in the CRC cell lines and HeLa cells but were rarely observed in human plasma, despite all three datasets being annotated using the same human GO BP database. Together, these results show that proteoform barcodes provide a simple and intuitive framework for comparing the functional composition of different biological samples.

### 3.2. Proteoform barcodes reveal different stages of breast cancer

We used our recently published human plasma TDP dataset of different stages of breast cancer^29^ to create proteoform barcodes and evaluated the potential of using proteoform barcodes for clustering human plasma samples into different breast cancer stages. Before generating the proteoform barcodes, principal component analysis (PCA) was performed to assess differences among the four groups. As shown in **Figure 3A**, the Control and G1 groups were clearly separated from the G2 and Metastatic groups, whereas partial overlap was observed between the Control and G1 groups and between the G2 and Metastatic groups. **Figure 3B**, **Figure S1**, and **Figure S2** show a total of 39 proteoform barcodes representing the 39 human plasma samples (9 Controls; 10 samples for stages G1, G2, and metastatic breast cancer, respectively) analyzed in our recent work.

**Figure 3.**
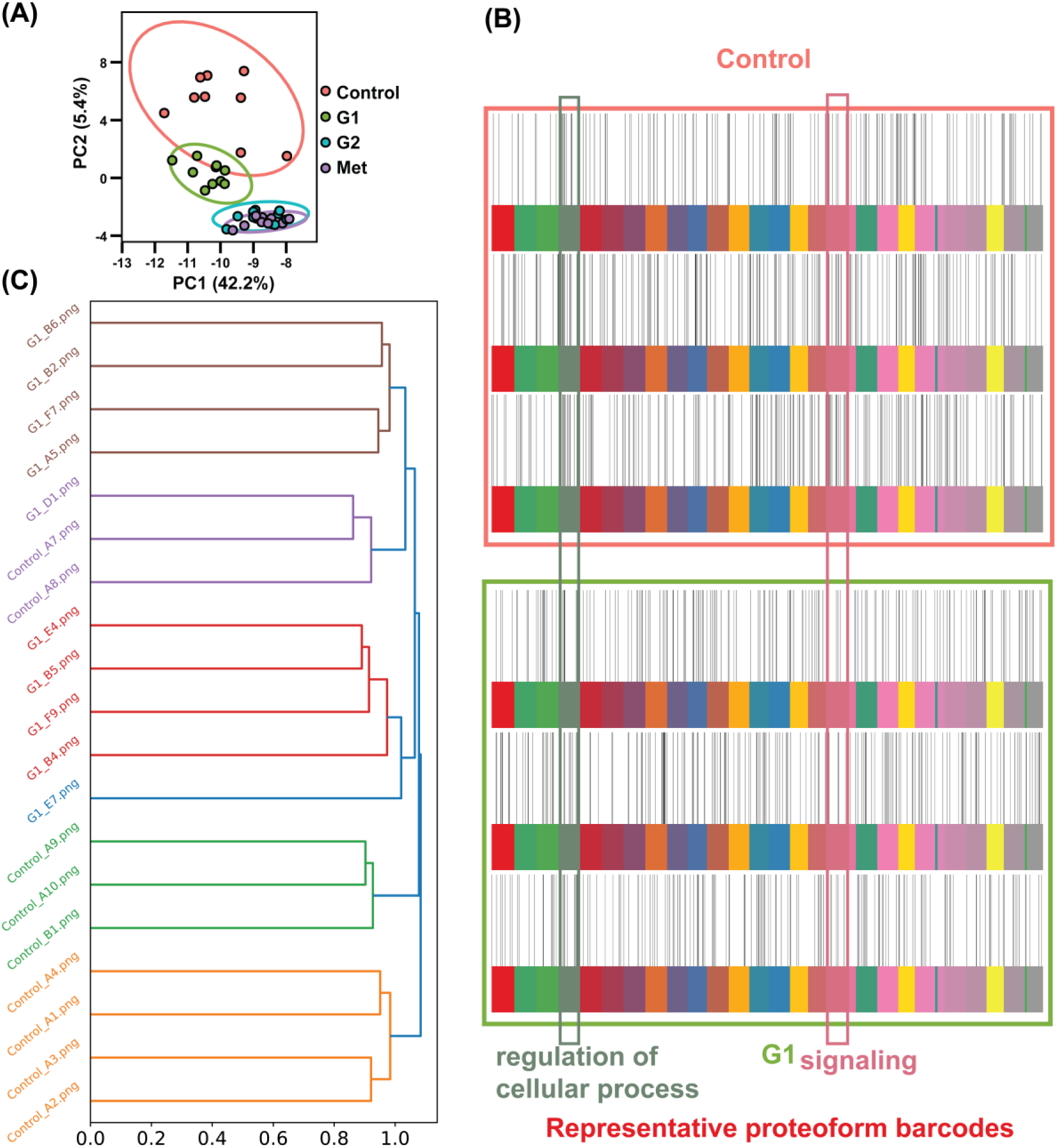
Summary of the human plasma proteoform barcode data. (A) PCA plot of plasma proteoforms from the control group and the three breast cancer stages [G1, G2, and Metastatic (Met)]. (B) Representative proteoform barcodes of the control and G1 groups constructed from all 3,785 proteoforms associated with the top 30 biological processes. (C) Hierarchical clustering of proteoform barcodes (Control and G1-stage breast cancer) using correlation distance and average linkage. Original sample names (e.g., G1_B6) were retained to facilitate tracking across analyses, and the same naming convention is used in all subsequent figures.

Compared with the Control and G1 groups (**Figure S1**), the G2 and Metastatic groups exhibited lower relative intensities for proteoforms associated with cellular process, primary metabolic process, and macromolecule metabolic process, **Figure S2**, suggesting that reduced abundance of proteoforms involved in these biological processes may be associated with breast cancer progression. In addition, the primary difference between the Control and G1 groups was observed in the regulation of cellular process and signaling (the green and pink BPs in **Figure 3B** and **Figure S1**), where fewer proteoforms were detected in G1 than in the Control group. Overall, the proteoform barcodes enabled clear visual discrimination between early-stage (Control/G1) and late-stage (G2/Metastatic) breast cancer. However, further studies with larger cohorts are required to evaluate their potential for early breast cancer diagnosis.

To evaluate whether proteoform barcode images capture breast-cancer-stage-associated differences, hierarchical clustering was performed using RGB intensity profile features extracted from the proteoform barcode images. Control and G1 barcodes were largely clustered according to their biological groups, although some Control samples were grouped with the G1 cluster, **Figure 3C**. This clustering pattern is consistent with the overlap observed between Control and G1 samples in the PCA plot (**Figure 3A**), suggesting that plasma proteoforms can be similar during the early stage of breast cancer development. Great clustering was observed for the Control versus G2, Control versus Metastatic, and G1 versus G2 comparisons (**Figure S3B–D**), consistent with the distributions in the PCA plot (**Figure 3A**). In contrast, G2 and Metastatic barcodes exhibited the least distinct clustering pattern (**Figure S3E**), suggesting that plasma proteoform profiles become increasingly similar during later stages of breast cancer progression. When all four groups were clustered simultaneously, G2 and Metastatic samples were predominantly grouped within the same major branch, whereas Control and G1 samples formed another major branch (**Figure S3A**). These results suggest that distinguishing early-stage breast cancer (G1) from Controls is more challenging than distinguishing later disease stages. Nevertheless, proteoform barcode clustering still captured important information across breast cancer stages.

Although the PCA score plot showed slight overlap between the Control and G1 groups, an overall separation trend was observed. To further improve the separation of G1 and Control groups for better early detection of breast cancer, we employed a machine learning approach, the random forest classifier, which used 60% of the samples for training and the remaining 40% as an independent test set and correctly classified all samples in the independent test set, achieving an accuracy of 100%. Stratified 5-fold cross-validation further yielded a mean ROC AUC of 1.000, indicating excellent and robust discrimination between Control and G1 samples, **Figure 4B**. However, these results should be interpreted with caution because the model was trained and evaluated using only 19 biological replicates. The limited sample size increases the risk of overfitting. Proteoform datasets from much larger cohorts will be used in our future studies.

**Figure 4.**
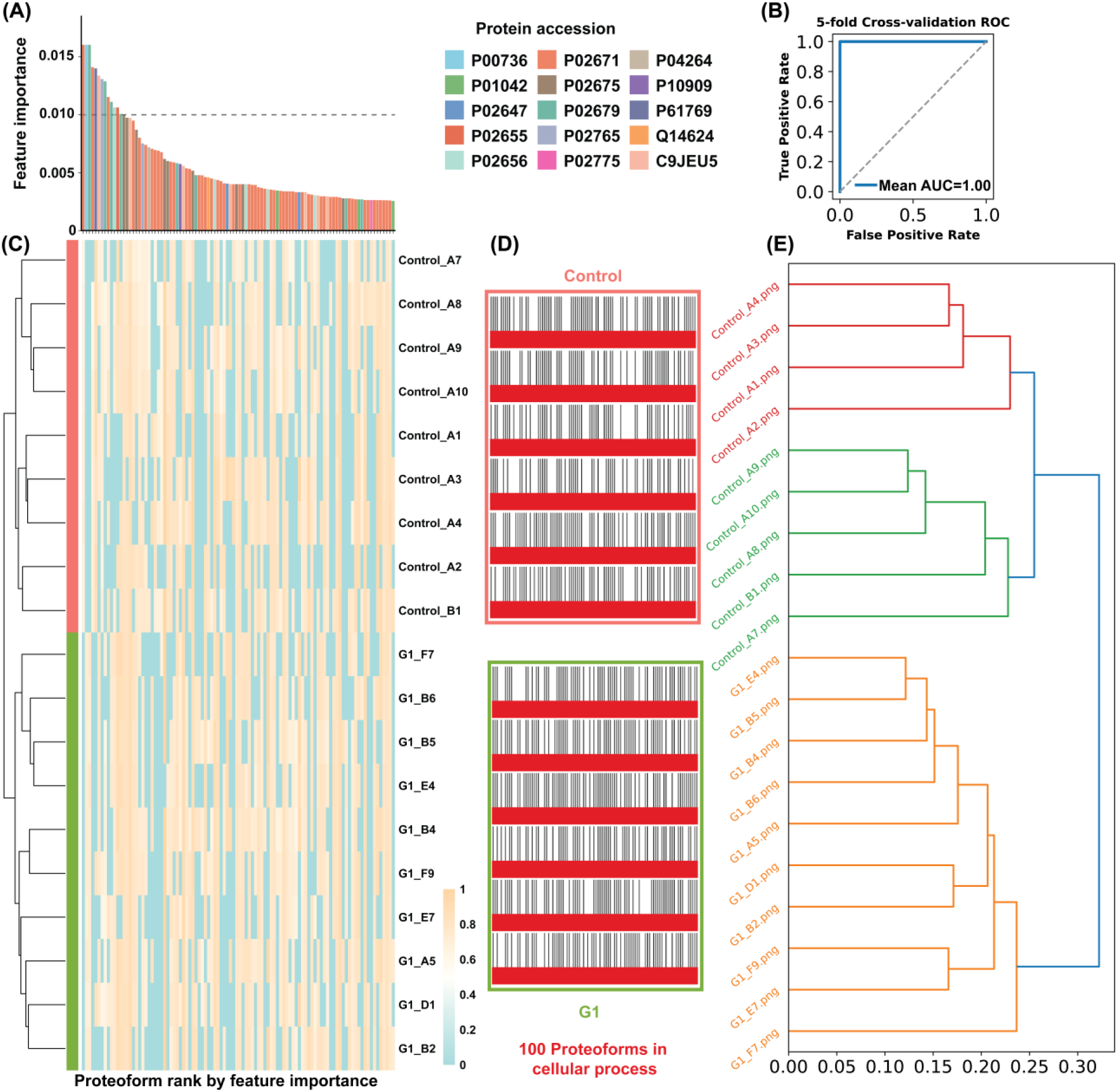
Improved proteoform barcodes of human plasma proteoforms for separating Controls and G1-stage breast cancer. (A) Top 100 proteoforms ranked by feature importance. Protein UniProt accession numbers are labeled on the plot. The x-axis is the same as in panel (C). (B) Five-fold cross-validation of the random forest classification model. (C) Heatmap of the top 100 proteoforms ranked by feature importance for distinguishing Control and G1 samples. Proteoforms are ordered according to their feature-importance ranks (1–100) with detailed proteoform information provided in Supporting Information II Sheet 17, and samples are hierarchically clustered based on normalized proteoform intensity using Euclidean distance and complete linkage. The color scale represents the normalized proteoform intensity. (D) Representative proteoform barcodes generated using the top 100 proteoforms selected by the random forest model. Control samples are outlined in pink, and G1 samples are outlined in green. (E) Hierarchical clustering of proteoform barcodes generated from the top 100 proteoforms using correlation distance and average linkage.

After confirming that the random forest model was capable of distinguishing Controls from G1 samples, the top 100 proteoforms ranked by feature importance were selected for heatmap visualization, among which 14 proteoforms had feature importance values greater than 0.01, **Figures 4A** and **4C**. The clustering pattern in the heat map further indicated that the selected proteoforms captured the differences between the two groups. In **Figure 4A**, proteoforms derived from the same protein are highlighted using the same color. Clusterin (CLU, P10909) has been reported as a biomarker of breast cancer prognosis^51, 52^. ITIH4 (Q14624)-derived peptides have been reported as potential biomarkers for breast cancer^53^. In addition, a plasma fragment of fibrinogen alpha chain (FGA, P02671), was identified as a potential biomarker for HER2-positive breast cancer and was reported to return toward normal levels after surgery^54^. Downregulated APOA1 (P02647) expression has also been associated with shorter overall survival in patients with breast cancer^55^.

Next, the top 100 proteoforms ranked by feature importance were selected to construct new proteoform barcodes, with the aim of making Control and G1 samples easier to distinguish by visual inspection. Interestingly, all 100 selected proteoforms were associated with the BP, cellular process. Therefore, the new proteoform barcodes were constructed using only the cellular process BP, in which each proteoform appeared only once in each barcode. As shown in **Figure 4D** and **Figure S4**, the new proteoform barcodes exhibited much clearer visual differences between the Control and G1 groups. The differences between the proteoform barcodes of the Control and G1 groups were readily distinguishable by visual inspection. To further evaluate whether the barcodes generated from the top 100 proteoforms could be more readily distinguished computationally than those generated from all proteoforms, hierarchical clustering was performed. The samples were separated into two major clusters, corresponding to the Control and G1 groups, indicating distinct proteoform barcode patterns between the two groups, **Figure 4E**. These results suggest that feature-selected proteoform barcodes provide a more discriminative representation of disease-associated proteoform patterns, making them well suited for computational analysis. We expect the panel of proteoforms represented as the feature-selected proteoform barcodes to be used as new biomarkers for early detection of breast cancer with the assistance of machine learning approaches. We also expect the concept of proteoform barcodes assisted by machine learning could become a standard approach for distinguishing different biological conditions, i.e., early detection of various diseases.

## 4. Conclusions

We provide the “proteoform barcode” as an intuitive visualization framework for TDP. Proteoform barcode allows a bird’s-eye view of proteoforms in biological samples and their associated biological processes. It converts a list of proteoforms to an image containing abundance and functional information, which could be used to represent specific sample species and to facilitate early diagnosis of diseases (i.e., cancer) assisted by machine learning/AI tools. Because proteoform barcodes transform complex proteomics data into standardized images, they can be directly analyzed using image-based AI. Even simple image analysis methods, such as hierarchical clustering, successfully captured biologically meaningful differences among breast cancer stages, demonstrating the feasibility of the approach. With larger clinical cohorts and more advanced AI models, proteoform barcodes have the potential to transform TDP in early disease diagnosis. In this study, we used the proteoforms identified by the TopPIC software^12^ to demonstrate the idea of “proteoform barcode”. Database search results from other TDP software [i.e., ProSight^13^, Informed-Proteomics^14^, MASH Suite Pro^15^, FLASHDeconv^16^, ProteoformX (https://www.bioinfor.com/proteoformx/), and Proteoform Suite^56^ can be easily modified to fit our procedure of producing proteoform barcodes. The idea of proteoform barcode could also be expanded to other omics data (i.e., bottom-up proteomics and metabolomics). We expect the proteoform barcode could become a general approach for data visualization in omics studies in the near future.

## Acknowledgments

We thank the National Institute of General Medical Sciences through the grant R35GM153479, and the National Cancer Institute (NCI) through the grant R01CA247863 for their support.

## Author contributions

Yifan Yue: Methodology, Software, Formal analysis, Visualization, Writing – original draft. Guangyao Gao, Fei Fang, and Guijie Zhu: Data curation. Seyed Amirhossein Sadeghi and Reyhane Tabatabaeian Nimavard: Resources. Liangliang Sun: Conceptualization, Methodology, Supervision, Writing – review & editing

