## Supporting Information I for "Proteoform Barcode: An Intuitive Visualization Framework for Top-Down Proteomics"

* Corresponding author

**Table S1**. Example proteoforms identified in previous top-down proteomics (TDP) studies.

| | Database protein sequence | Adjusted precursor mass | Proteoform | Protein accession |
|---|---|---|---|---|
| 1 | SRKESYSVYVYKVLKQVHPDTGISSKAMGIMNSFVNDIFERIAGEASRLAHYNKR | 6302.42 | R.SRKESYSVYVYKVLKQVHPDTGISS(KAM)[13.1882]GIMNSFVNDIFERIAGEASRLAHYNKR.S | sp\|O60814\|H2B1K_HUMAN |
| 2 | SRKESYSVYVYKVLKQVHPDTGISSKAMGIMNSFVNDIFERIAGEASRLAHYNKR | 6305.224 | R.SRKESYSVYVYKVLKQVHPDTGISSKAMG(I)[15.9920]MNSFVNDIFERIAGEASRLAHYNKR.S | sp\|O60814\|H2B1K_HUMAN |
| 3 | GLTLHLKFLEPFDIDDHQQVHCPYDQLQIYANGKNIGEFCGKQRPPDLDTSSNAVDLLFFTDESGDSR | 7704.65293 | R.GLTLHLKFLE(PFDIDDHQQVHCPYDQLQIYANGKNIGEFCGKQRPPDLDTSSNA)[-2.0258]VDLLFFTDESGDSR.G | sp\|P00736\|C1R_HUMAN |
| 4 | GLTLHLKFLEPFDIDDHQQVHCPYDQLQIYANGKNIGEFCGKQRPPDLDTSSNAVDLLFFTDESGDSR | 7704.65737 | R.GLTLHLKFLEPFDID(DHQQVHCPYDQLQIYANGKNIGEFCGKQRPPD)[-2.0214]LDTSSNAVDLLFFTDESGDSR.G | sp\|P00736\|C1R_HUMAN |

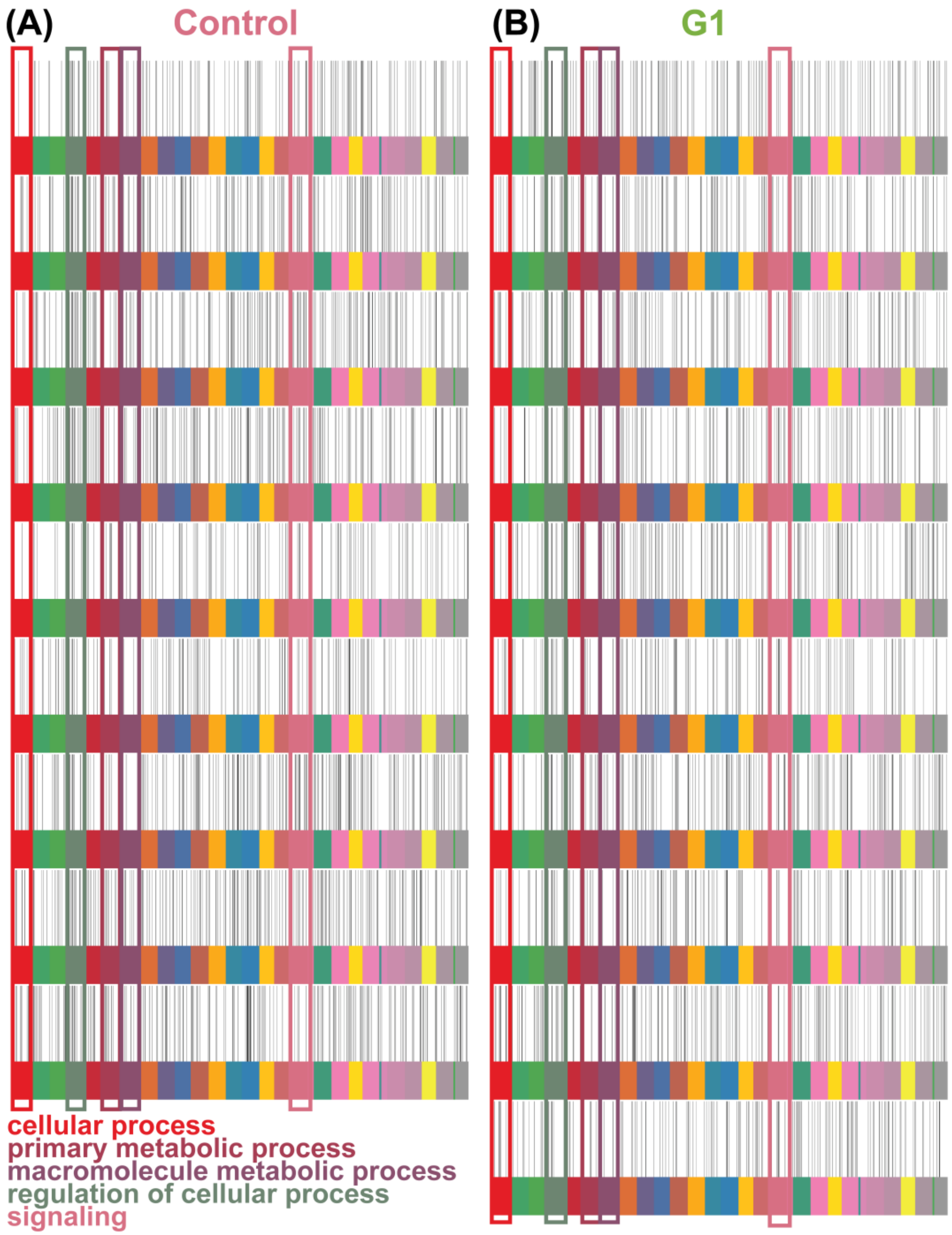


**Figure S1.** Proteoform barcodes of Control and G1 breast cancer plasma samples constructed using all proteoforms associated with the top 30 GO Biological Process terms. Each colored band at the bottom represents a different biological process (BP) and contains multiple vertical lines, with each vertical line representing a proteoform associated with that BP. Because a proteoform may be associated with multiple BPs, the same proteoform may appear in multiple colored bands. The grayscale gradient of the vertical lines represents normalized intensity of each proteoform, ranging from 0 (white) to 1 (black); the same scale is used in all subsequent figures.

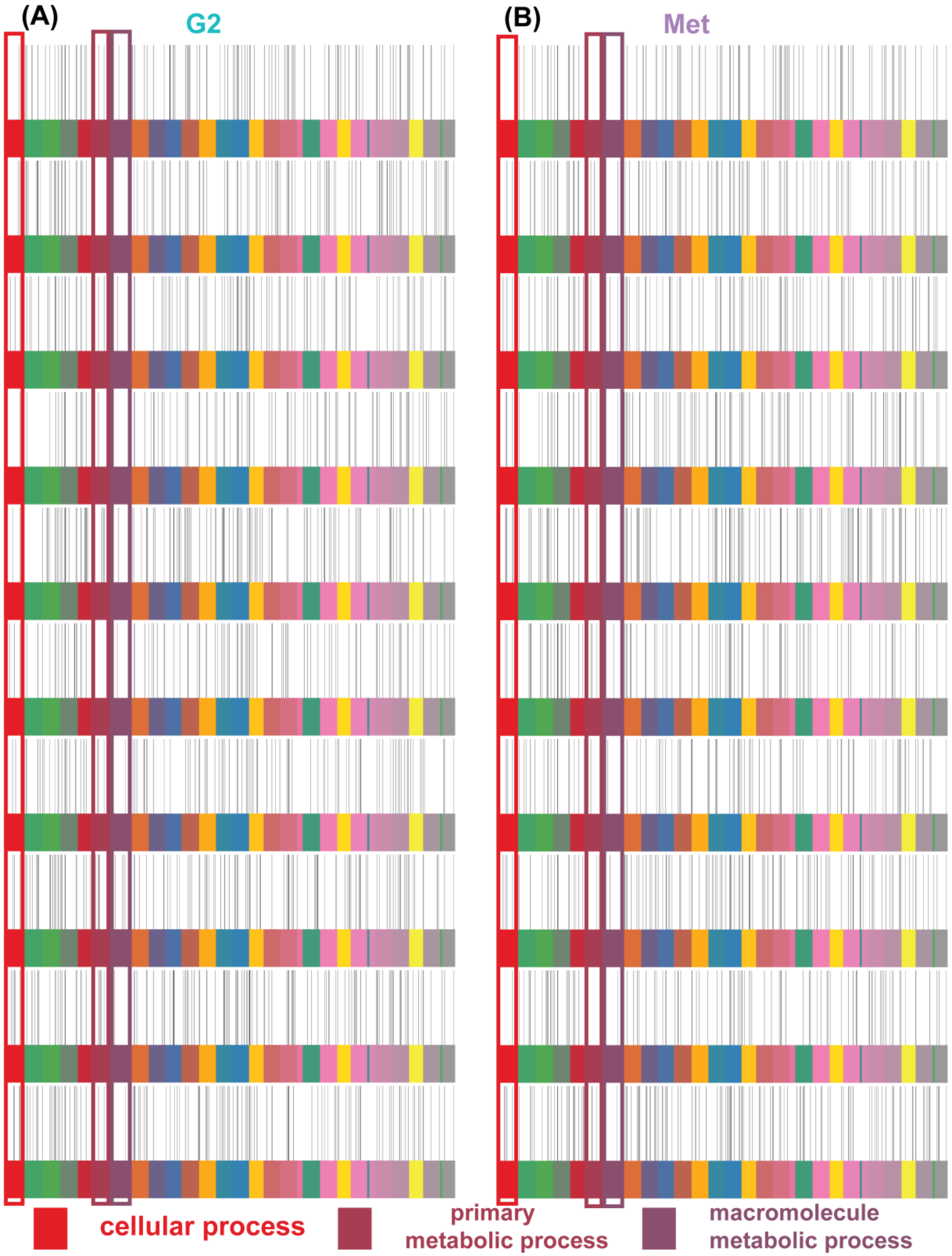


**Figure S2.** Proteoform barcodes of G2 and metastatic breast cancer plasma samples constructed using all proteoforms associated with the top 30 GO Biological Process terms.

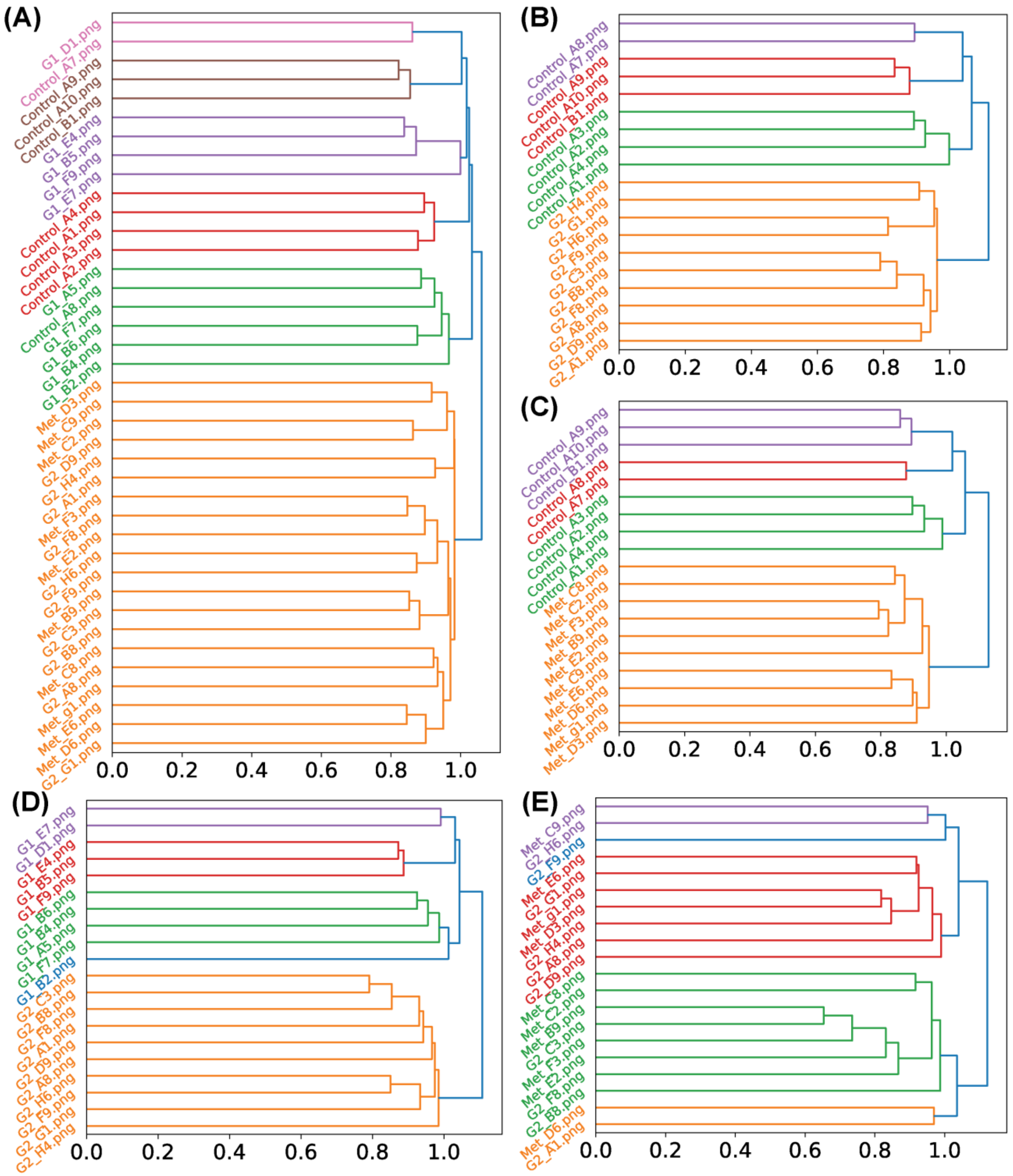


**Figure S3.** Hierarchical clustering of proteoform barcode images. (A) All experimental groups. (B) Control vs. G1. (C) Control vs. Metastatic. (D) G1 vs. G2. (E) G2 vs. Metastatic. Original sample names (e.g., G1_B6) were retained to facilitate tracking across analyses.

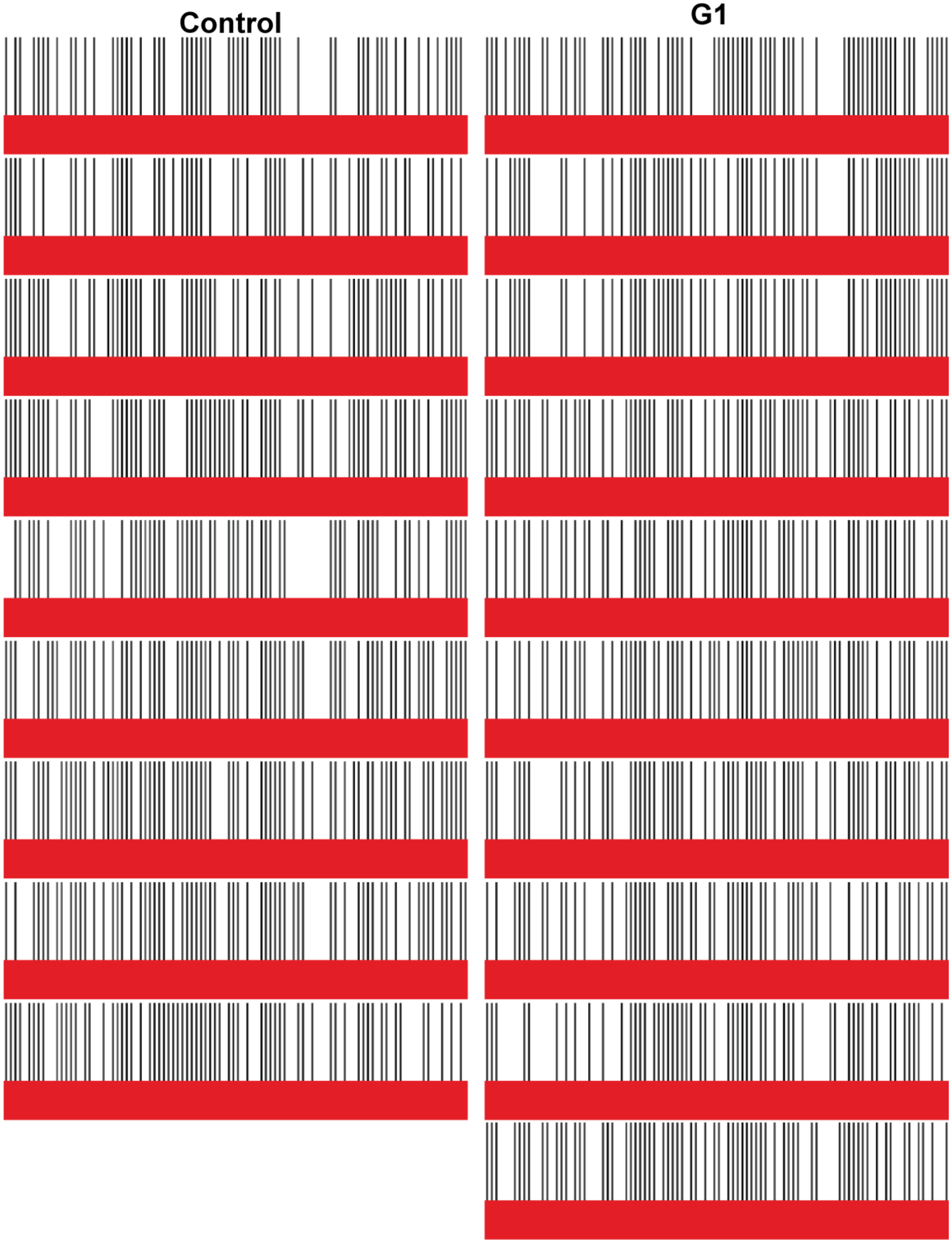


**Figure S4.** Proteoform barcodes of Control and G1 breast cancer plasma samples constructed using the top 100 proteoforms selected by random forest feature importance. All selected proteoforms were associated with the cellular process BP term.
